# An ectosymbiosis-based mechanism of eukaryogenesis

**DOI:** 10.1101/2020.03.25.009100

**Authors:** Tze-Fei Wong, Chung-Kwon Chan, Xi Long, Hong Xue

**Affiliations:** Division of Life Science, Hong Kong University of Science and Technology, Hong Kong, China

**Keywords:** *Aciduliprofundum*, archaeal parent, ectosymbiosis, endosymbiosis, eukaryogenesis, valyl-tRNA synthetase

## Abstract

The mechanisms proposed for eukaryogenesis are divisible into mitochondria-early and mitochondria-late ones, where the mitochondriate-eukaryotes (MTEs) were evolutionary precursors or products of the amitochondriate-eukaryotes (AMIs) respectively. Analysis of prokaryote-to-eukaryote gene transfers in eukaryogenesis showed two tranches of high-intensity transfers from prokaryotes to eukaryotes mediated by endosymbioses that gave rise to mitochondria and chloroplasts, and hundreds of medium-intensity transfers which included the transfer of hydrogenase and pyruvate: ferredoxin oxidoreductase genes from the *Thermoanaerobacter-Hungateiclostridium-Sporanaerobacter* group to the AMIs. Since 94.5% of these medium-intensity transfers generated more than 100 inter-proteome similarity hits between each donor-recipient pair, they were not readily explicable by horizontal gene transfers (HGTs) or endosymbioses, pointing instead to the participation of a large number of ectosymbiotic transfers. The euryarchaeon *Aciduliprofundum boonei* (Abo) and the gammaproteobacterium *Escherichia coli* (Eco) were the foremost contributors of archaeal and bacterial genes for the eukaryotic DNA-apparatus respectively, and the Abo/Eco ratios for the prokaryote-derived genes in eukaryotes decreased from Microsporidia to other AMIs, and even more so in the MTEs. These findings supported an ectosymbiosis-based mechanism of eukaryogenesis with Abo as the Archaeal Parent of Eukarya, and Microsporidia as the eukaryotes phylogenetically closest to the Last Eukaryotic Common Ancestor (LECA).

## INTRODUCTION

For over a century endosymbiosis has been regarded as a core participant in eukaryogenesis^1^, with the development of the eukaryotic cell being hosted within an archaeal parent^2,3^; within the Bacteria domain in the three-domain structure of life^4^; starting from a chimeric fusion of bacteria and archaebacteria with a single nucleus, a single kinetosome, and protein connectors^5^; from the Archezoa such as a phagotrophic archezoan similar to *Mastigamoeba*^6^; and within a hydrogen-dependent host^7^ along with other formulations.

Different lines of evidence have favored an Archaeal Parent for the Eukarya domain: *Aciduliprofundum boonei* (or Abo; see three-letter abbreviations in Table 1) furnished a well-endowed candidate Archaeal Parent with top inter-proteome bitscores toward the AMIs *Giardia* and *Trichomonas*^8^; the Asgard and TACK archaeons provided important genes for eukaryotic signature proteins^9^ to the eukaryots^10,11^; a large excess of archaea-derived over bacteria-derived ribosomal proteins was found in *Giardia, Trichomonas*, yeast, and humans^8^; and archaeal genes were more important than bacterial genes for the eukaryotes^12^. However, the question of whether the MTEs emerged early prior to the AMIs, or late following the AMIs has to be resolved. Although studies on SSU rRNA, elongation factor EF-1alpha and other proteins favored older ages of AMIs relative to MTEs^13-16^, the discoveries of mitochondrial genes in the AMIs have gained momentum for the *degeneration theory* that AMIs arose from the degeneration of MTEs, thereby favoring older ages of MTEs in comparison with AMIs^17-22^. Recently, a compromise between these views has also been introduced to the effect that, if observations were not completely compatible with a Last Eukaryotic Common Ancestor (LECA) with upfront mitochondria, the formation of some Pre-endosymbiont^23^ or First Eukaryotic Common Ancestor (FECA)^10^ might render possible a more gradual initiation of eukaryogenesis.

**Table 1.**
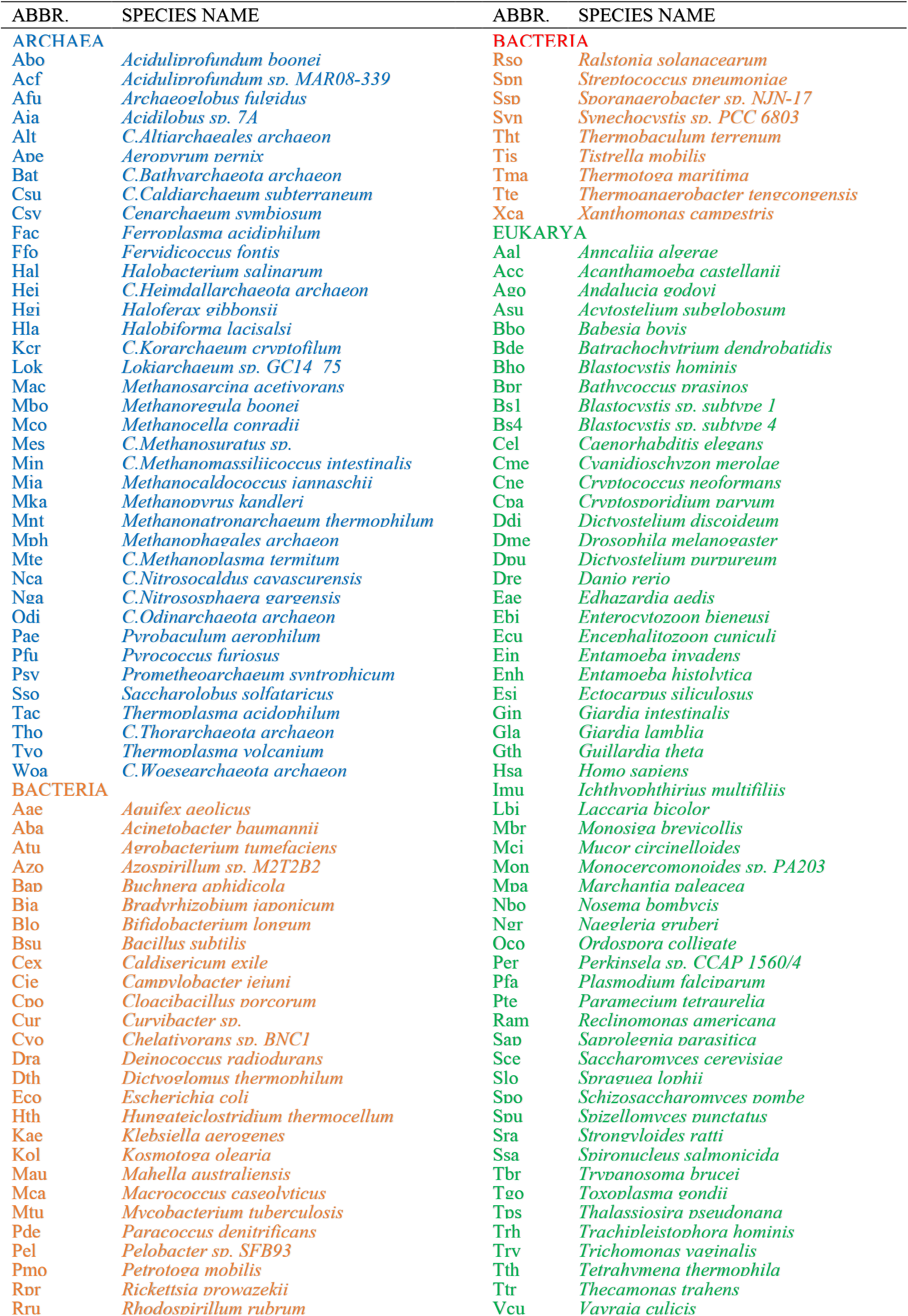
Species names and their three-letter abbreviations. See Supplementary Table 1 for descriptions of species.

Earlier, examination of the contents of prokaryote-derived proteins in Gla and Trv demonstrated that, while the levels of archaea-derived proteins were comparable in Gla and Trv, bacteria-derived proteins were much higher in Trv compared to Gla^8^. Since the influx of these exogenous proteins could be related to the development of LECA, Pre-endosymbiont, or FECA, the present study was directed to an analysis of their biological sources, avenues of entry into eukaryotes, and whether they represented a boon or hindrance to eukaryogenesis. The analysis led to the establishment of Abo as the Archaeal Parent of eukaryotes, the location of LECA within or near Microsporidia, and an ectosymbiosis-based mechanism of eukaryogenesis.

## METHODS

### Source of sequences

VARS sequences from different species were retrieved mostly from NCBI GenBank release (ftp://ftp.ncbi.nlm.nih.gov/genomes/)^24^. Mitochondrial DNA-encoded protein sequences were retrieved from the RefSeq mitochondrial reference genomes in the NCBI Protein database (https://www.ncbi.nlm.nih.gov/protein)^25^.

### Estimation of inter-proteome or inter-protein similarity bitscores

Inter-proteome and inter-protein similarity bitscores were performed as described^8^. The proteomes of various species were employed to construct a local BLAST database using makeblastdb^26^, and query proteomes or proteins were searched against the local database using BLASTP with a BLOSUM62 matrix and thresholds set to e-value <1×10^−5^, >25% percent identity and >50% query coverage. Only when the query and subject sequences that were the best match of each other, viz. when query sequence *n* from species 1 exhibited the highest bitscore toward subject sequence *m* among all proteins of species 2 and vice versa, were the data included in the estimation of similarity.

## RESULTS

### Heatmap of prokaryote-to-eukaryote gene transfers

To survey the gene transfers from prokaryotes to eukaryotes in the course of eukaryogenesis, a heatmap of the similarity bitscores between different prokaryote (on the x-axis) and eukaryote (on the y-axis) protein-coding genes was constructed. It showed numerous plausible transfers of genes from prokaryotes to eukaryotes at various x-y junctions. The large transfers from proteobacteria to the MTEs, and from Syn (representing cyanobacteria) to the algae Cme, Esi, Gth, Tps, and Bpr would correspond to the genes originating from the mitochondria- and chloroplast-yielding bacterial endosymbionts respectively. The transfer of Syn genes mainly to the algae with only limited spillover to other eukaryotes attested to the high specificity of the similarity bitscores in identifying cognate pairs of gene donors and recipients. There were a wide variety of medium-intensity transfers from archaea and bacteria into both the AMI and MTE eukaryotes (Figure 1). These medium-intensity transfers were unlikely to be the outcome of HGTs, because 94.5% of them consisted of more than one hundred similarity hits each (Supplementary Table S2), whereas HGTs usually bring about transfers of single genes. Instead, that they consisted largely of ectosymbiotic gene transfers was suggested by the parallel usage of endosymbiosis as in legume-*Rhizobium* interactions, and ectosymbiosis as in the binding of *Nostoc* to specialized leaf cavities on *Anthoceros* host in nitrogen fixation^27^; the critical role of ectosymbiosis in determining benthic biodiversity in the Arctic deep sea^28^; protist-spirochete interactions in the termite gut^29^; the exchanges of genes between DPANN and *Thermoplasma*^30^; and the repeatedly evolved host-specific ectosymbiosis between amphipods and sulfur-oxidizing bacteria in a cave ecosystem^31^. As well the results in the heatmap were in accord with the findings of a wide variety of archaeal and bacterial genes in the genome of yeast^32^ as well as Gla and Trv^8^, and the apparent lack of any Pre-endosymbiont, FECA, or endosymbionts aside from the proteobacterial and cyanobacterial ones that led to the formation of mitochondria and chloroplast respectively^33^.

**Figure 1.**
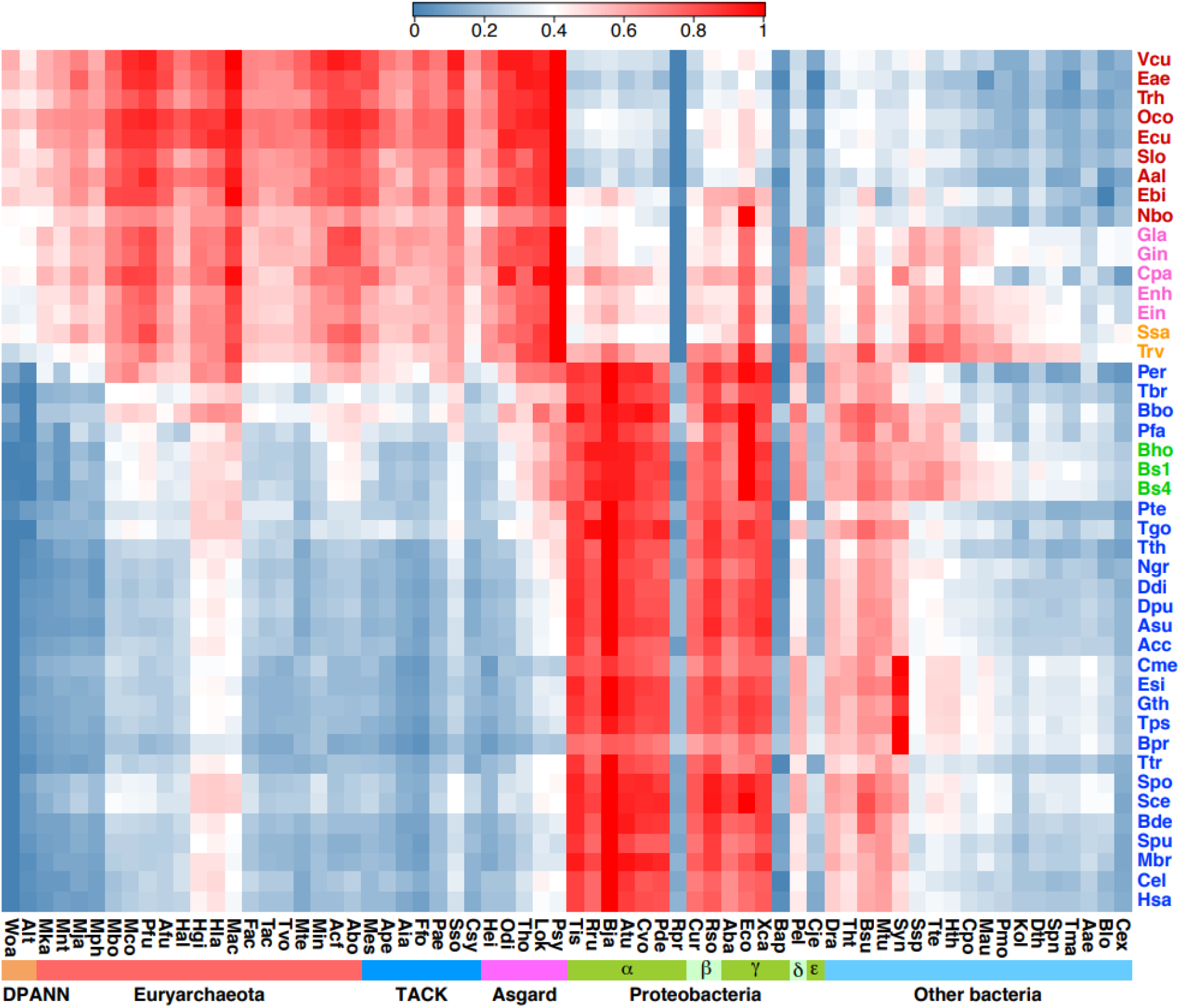
Heatmap of inter-proteome similarity bitscores between eukaryotes and prokaryotes. For each eukaryotic proteome, its similar bitscore toward a prokaryotic proteome is represented by the square at the intersection between a eukaryotic row and a prokaryotic column, and scaled linearly from 0 to 1.0 for each row according to the thermal scale, with zero marking the minimum and 1.0 the maximum bitscores. On the y-axis, AMIs with mitosomes are placed at the top followed by AMIs with hydrogenosomes and MTEs. Different MTEs are ordered on the y-axis broadly by biological groups and according to the Abo/Eco ratios for the different groups (Figure 7). The aggregate similarity bitscores for individual prokaryote-eukaryote pairs are shown in Supplementary Table 2.

Moreover, the display by some archaea of an *accelerated gene adoption* (AGA) phenotype that enriched their genomes with more bacteria-derived genes than other archaeons indicated that the highly AGA-active archaeons could recruit bacterial genes efficiently through non-HGT or hyper-HGT mechanisms, and few of the genes belonging to a subset of bacteria exemplified by Rpr, Bap, Cje, and Blo resisted recruitment via AGA^8^. Likewise, on the heatmap in Figure 1, few Rpr, Bap, Cje and Blo genes were transferred to any AMI or MTE archaeons, suggesting that the Archaeal Parent excelled, like Abo or Tvo, in AGA-activity, making it a superb recruiter of ectosymbiosis-transferred prokaryotic genes in the course of eukaryogenesis.

### Identification of Archaeal Parent

Since it was likely that the earliest eukaryotes inherited some elements of the information system of their prokaryote ancestor, the eukaryotic DNA-apparatus would be an attractive site to look for such ancestral footprints. Accordingly, the similarity bitscores exhibited by a range of prokaryotes toward the DNA-apparatus genes, viz. Cluster of Ortholog Groups^34^ for replication, recombination and repair (COG-RRR), and for replication and repair (COG-L), of a variety of eukaryotes were estimated (Figure 2). The results demonstrated that:

**Figure 2.**
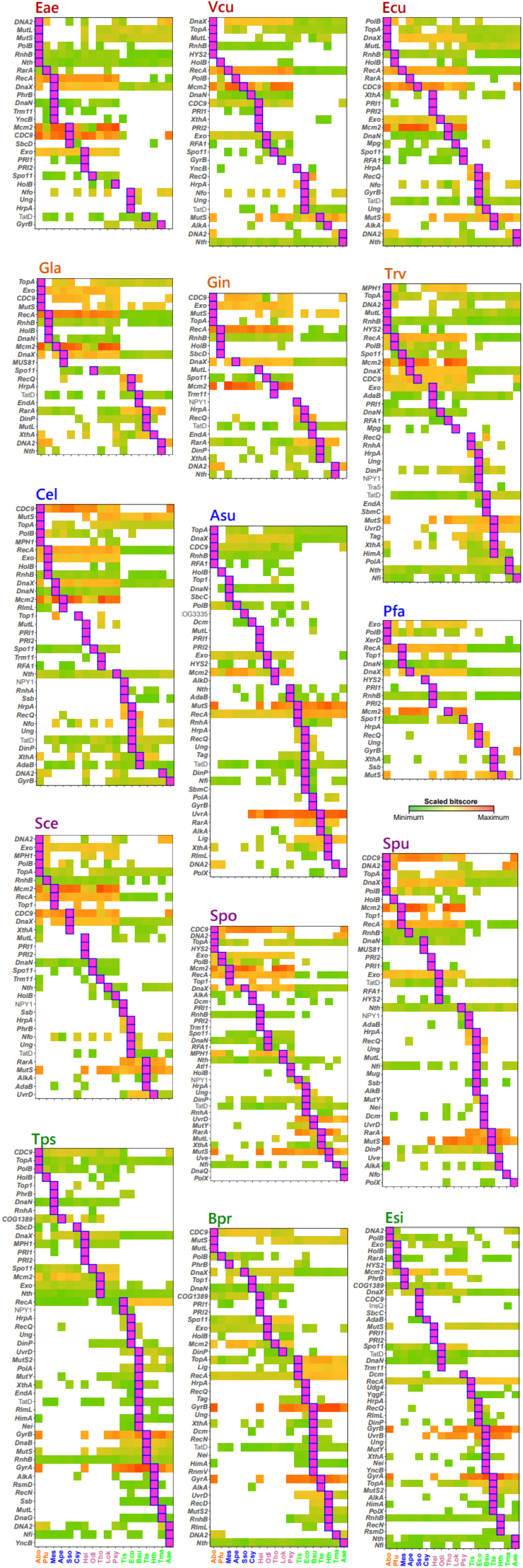
Similarity bitscores of COG groups in the eukaryotic DNA apparatus. Different eukaryote panels are divided into Microsporidia (labeled red), Excavata (orange), mixed MTEs (blue), Fungi (purple) and Algae (green). Similarity bitscores displayed toward various COG groups (y-axis) by different prokaryotic origins of genes (x-axis) are color-coded according to the thermal scale. Within each row, a purple rectangle inside a blue box marks the top bitscore for the row.

a. A sizable number of archaeons and bacteria exhibited top similarity bitscores toward the DNA apparatus in a variety of eukaryotes, in accord with the indication by the heatmap that both archaea and bacteria donated protein-coding genes to the eukaryotes through ectosymbiosis.
b. Abo exhibited the largest number of top bitscores along with Eco toward the Microsporidia Eae and Vcu, the largest number of top bitscores toward the *Giardias* Gla and Gin, the *Trichomonas* Trv, and a number of top bitscores toward Fungi and *C. elegans* but few toward the algae, in support of an AMI-Fungus-Animal central lineage in eukaryotic evolution.
c. *B. subtilis* exhibited prominent or co-prominent bitscores toward the algae.
d. The Asgard and TACK archaeons exhibited relatively few top bitscores, except for *Methanosuratus* (Mes) which shared the largest exhibitor role with Abo toward Eae. Interestingly, while Abo was co-prominent with Mes as top bitscore exhibitor toward the microsporidian Eae, it was co-prominent with *Pyrococcus* (Pfu) as top bitscore exhibitor for *Giardia*, suggesting that the phylogenetic branching of AMIs between Microsporidia and Excavata was accompanied by divergent contents of prokaryotic proteins.

These results were indicative of the foremost contributions made by Abo to the DNA-apparatus proteins of eukaryotes, especially the AMIs, which supported Abo as a leading candidate Archaeal Parent, and the emergence of AMIs prior to MTEs. In comparison, because Eco and Bsu genes were continually being recruited into both AMIs and MTEs on the heatmap, their genes became more eminent than the Abo genes among non-fungal and non-animal MTEs.

### Mitochondrial DNA-encoded and mitochondria-like organelle proteins

While there was convincing evidence on the alphaproteobacterial origin of mitochondria^35^, the indication in the heatmap of large influxes of multiple proteobacterial proteins into the MTEs was consistent with the discoveries of chimerism among mitochondrial proteins^36-38^. A substantial fraction of mtDNA-encoded proteins showed top or high similarity bitscores toward the alphaproteobacterium *Tistrella* (Tis) (Figure 3 top panel), in agreement with the proximity between Tis DNA and mtDNA on the phylogenetic tree for alphaproteobacteria^39^. However, proteins from other alphaproteobacteria such as *Orientia, Pelagibacter* and *Paracaedibacter* also displayed top bitscores toward some species of mitochondria. For human mtDNA-encoded proteins HM1-11, Tis likewise shared top bitscores with *Rickettsia* and *Ehrlichia* (Figure 3 lower panel). Since *Rickettsia* and *Ehrlichia* are infectious agents of humans, their acquisitions of top bitscores for HM3 and HM2 respectively might have occurred during the infection process. In any case, the entry of multiple genomic sequences from different alphaproteobacteria into the same species of mitochondria was reminiscent of the competition between distinct strains of *Nostoc* ectosymbionts for binding to the leaf cavities of their *Anthoceros* host in nitrogen fixation^27^, suggesting that different alphaproteobacteria could gain access to the mtDNA as ectosymbionts, where they would insert their genes or gene fragments through recombination with the alphaproteobacterial DNA already positioned as the resident mtDNA.

**Figure 3.**
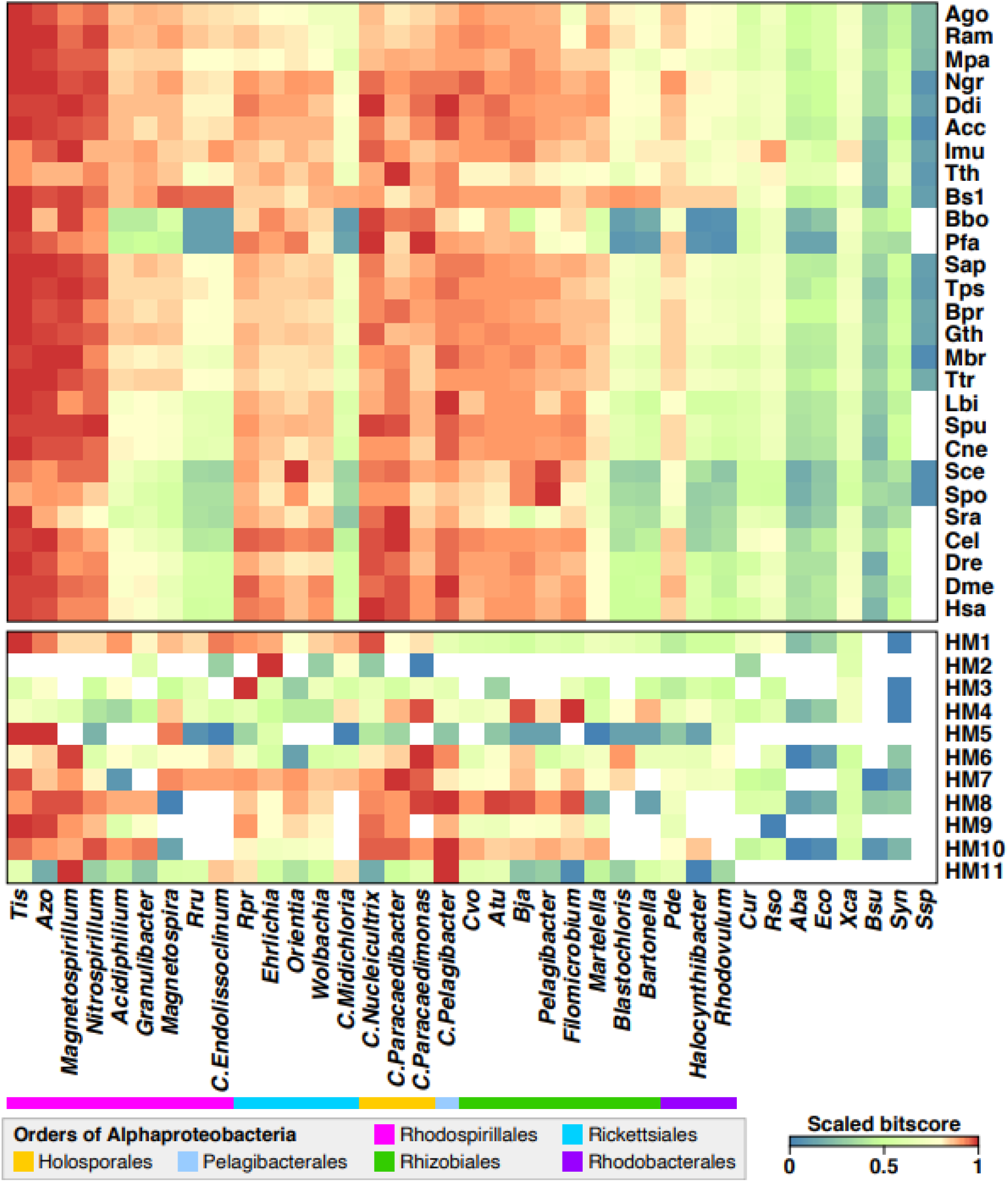
Bitscores of mitochondrial constituents. (Upper panel) Bitscores between mtDNA-encoded proteins of different eukaryotes and various bacterial proteomes. (Lower panel) Bitscores between human mtDNA-encoded proteins (HM1-11, viz. NADH dehydrogenase subunits 1-4, 4L and 5, cytochrome b, cytochrome oxidase subunits I-III, and ATP synthase F0 subunit-6 respectively) and various bacterial proteomes.

The origins of the hydrogenase and pyruvate:ferredoxin oxidoreductase (PFO) genes required for the activities of the mitosomes and hydrogenosomes of AMIs remained to be determined. Hydrogenosome resembled *Clostridium* metabolically^40^; and the PFOs from *Trichomonas, Giardia, Spironucleus*, and *Entamoeba* shared a single bacterial origin, even though the data were insufficient to identify that origin^41^. Since the heatmap pointed to medium-intensity ectosymbiotic gene transfers from the Clostridial/Firmicute species Tte, Hth and Ssp to Microsporidia, Gla, Gin, Enh, Ein, Ssa, Trv as well as MTEs like Bbo and Bho, the hydrogenase and PFO gene sequences from Tte were employed as specific probes for these genes, and the positive responses from AMIs and MTEs with mitosomes or hydrogenosomes (Figure 4, lines 9-10) suggest that the genes for their hydrogenase and PFO genes originated from bacteria related to the Tte-Hth-Ssp group; most of these species with positive responses were devoid of the genes for the subunits of the electron-transport proteins succinate dehydrogenase and fumarate reductase. In contrast, a variety of MTEs responded positively to the dehydrogenase and reductase probes from Pde (lines 11-12), in accord with their possession of an electron-transport chain^42^.

**Figure 4.**
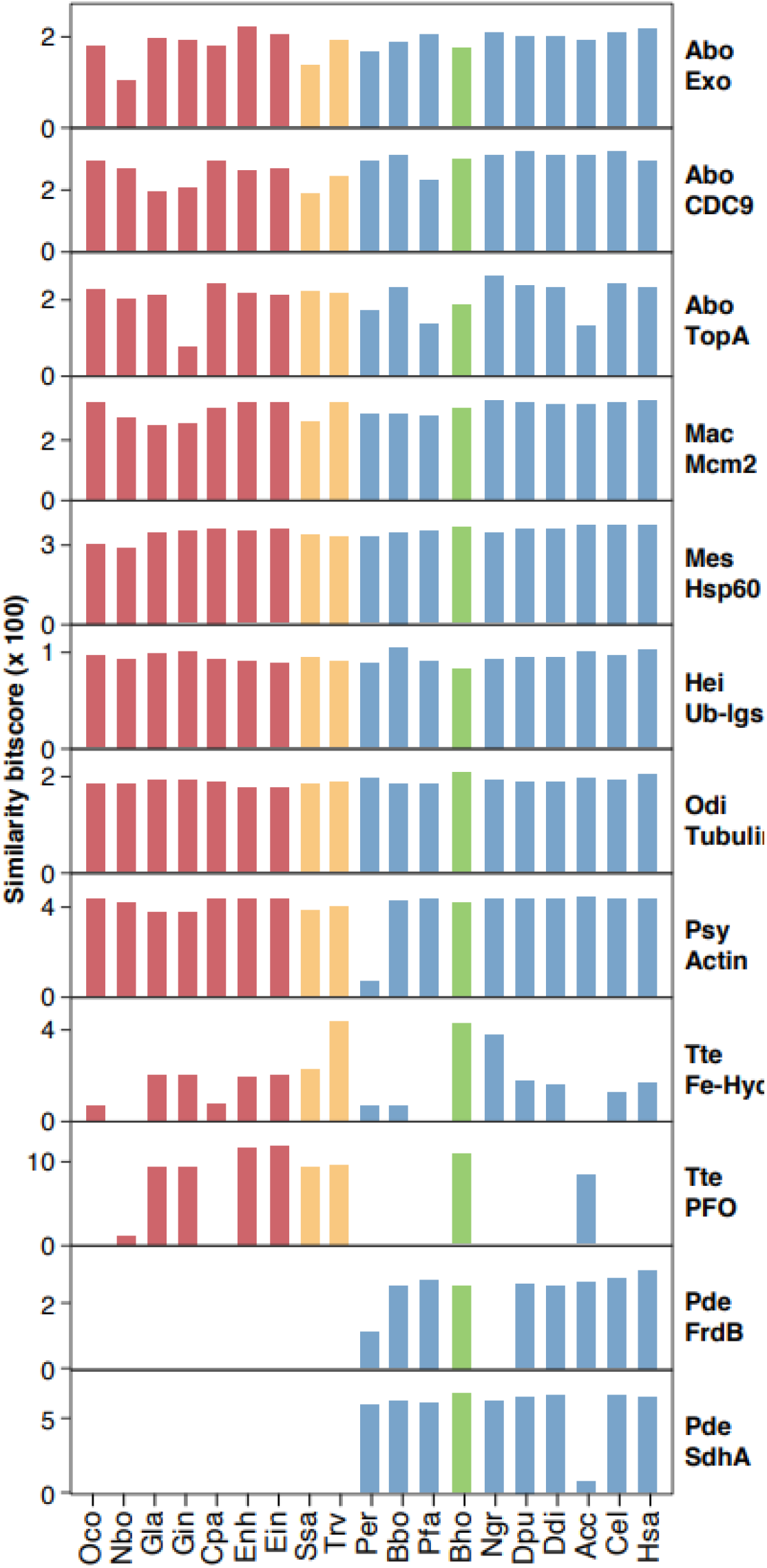
Similarity bitscores between prokaryotic probes and different eukaryotes. The bitscores pertained to probes based on Exo, 5’-3’ exonuclease; CDC9, ATP-dependent DNA ligase; TopA, DNA topoisomerase IA; Mcm2, DNA replicative helicase MCM subunit Mcm2; Ub-lgs, ubiquitin-protein ligase; Fe-Hyd, iron only hydrogenase; FrdB Fe-S protein subunit, and SdhA flavoprotein subunit, of succinate dehydrogenase/fumarate reductase. Lines 1-8 show fairly uniform responses from different eukaryotes to some non-electron transport protein probes, while lines 9-12 show the highly varied responses from distinct groups of eukaryotes to electron-transport protein probes.

### Degeneration theory of the origins of AMIs

Diplomonads, parabasalids and Microsporidia, all AMIs devoid of mitochondria, were postulated to be primitive eukaryotes that evolved prior to the advent of alphaproteobacteria-derived mitochondria based on the sequences of SSU rRNAs and protein markers such as elongation factor EF-1alpha^13-16^. However, this postulate was beset by possible long branch artefacts of rapidly evolving SSU rRNA, and an insert in EF-1alpha sequences, thereby favoring the degeneration theory that the AMIs were formed from degenerating Fungi^43,44^. For instance, the discovery of mitochondrial-like chaperonin 60 (cpn 60) genes in Trv, Gla and Microsporidia was regarded as evidence for the derivation of AMI VARS from MTE VARS^45^. However, the AMI and MTE VARS sequences were separated into distinct divisions on the cpn60 phylogenetic tree without clear indication of a mitochondrial origin of the VARS in AMIs. The application of the degeneration theory to the cpn60 genes was accordingly burdened with directional ambiguity, for these genes could have migrated of the protein from mitochondria to nucleus or *vice versa*^46^, as exemplified by the Arabidopsis mitochondrial genome which contained 16 sections of plastid DNA, fragments of nuclear genes, retrotransposons and sequences similar to plant-pathogen RNA viruses^47^.

In another application of the degeneration theory, the VARs sequences of Trv, Gla, a variety of MTEs and the bacteria Eco and *H. influenzae* were found to display a 37-aa insert just downstream from the KMSK motif, which was explained in terms of the derivation of Trv and Gla from degenerating MTEs^48^. However, the absence of this insert from all archaea ruled against its bequeathal to the eukaryotes by any Archaeal Parent; and the presence of an IKDENG insert in the VARS from Eco but not *Xanthomonas* (Xca) (Figure 5) pointed to an Xca rather than Eco origin of eukaryotic VARS. The Xca origin was supported by the conservation of amino acid residues (red asterisks) between the Xca and eukaryotic VARS in the vicinity of the KMSK motif. When a maximum parsimony phylogenetic tree rooted by Xca was built for the eukaryotic VARS sequences, it allocated the microsporidian VARS sequences to a distinct division on the tree, and Trv VARS to a particularly low-branching position near the root (Figure 6). These tree features were consistent with the ectosymbiotic transfer of the 37-aa insert from Xca to Trv, which in turn passed it to the microsporidian and non-microsporidian divisions on the tree. Within the non-microsporidian division, the two *Giardia* species were lower branching than the MTEs which included the Fungi. Therefore the application of the degeneration theory to the 37-aa insert in VARS was rendered uncertain by directional ambiguity, and erroneous in its supposition of the derivation of *Giardia* from degenerating Fungi.

**Figure 5.**
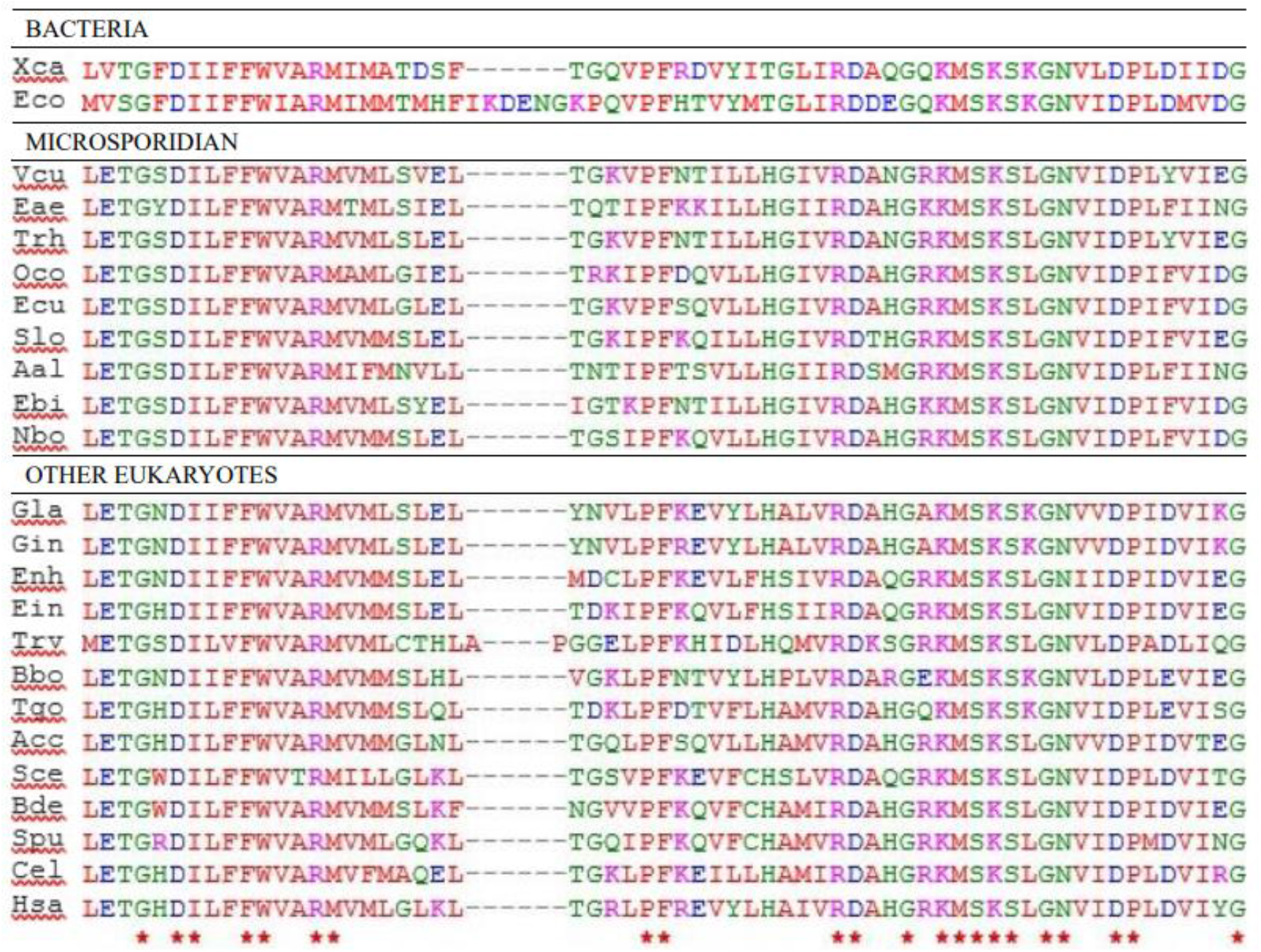
Alignment of VARS sequences bearing a 37-aa insert downstream from the KMSK motif. Amino acid residues that were totally conserved between all the aligned bacterial, microsporidial, and other eukaryotic VARS sequences are marked by red asterisks below the Hsa sequence. The aligned sequences shown are numbered according to the VARS sequence of Xca (see Supplementary Figure 1), starting from residue 699 and ending with residue 765. The KMSK motif is located at residues 748 – 751.

**Figure 6.**
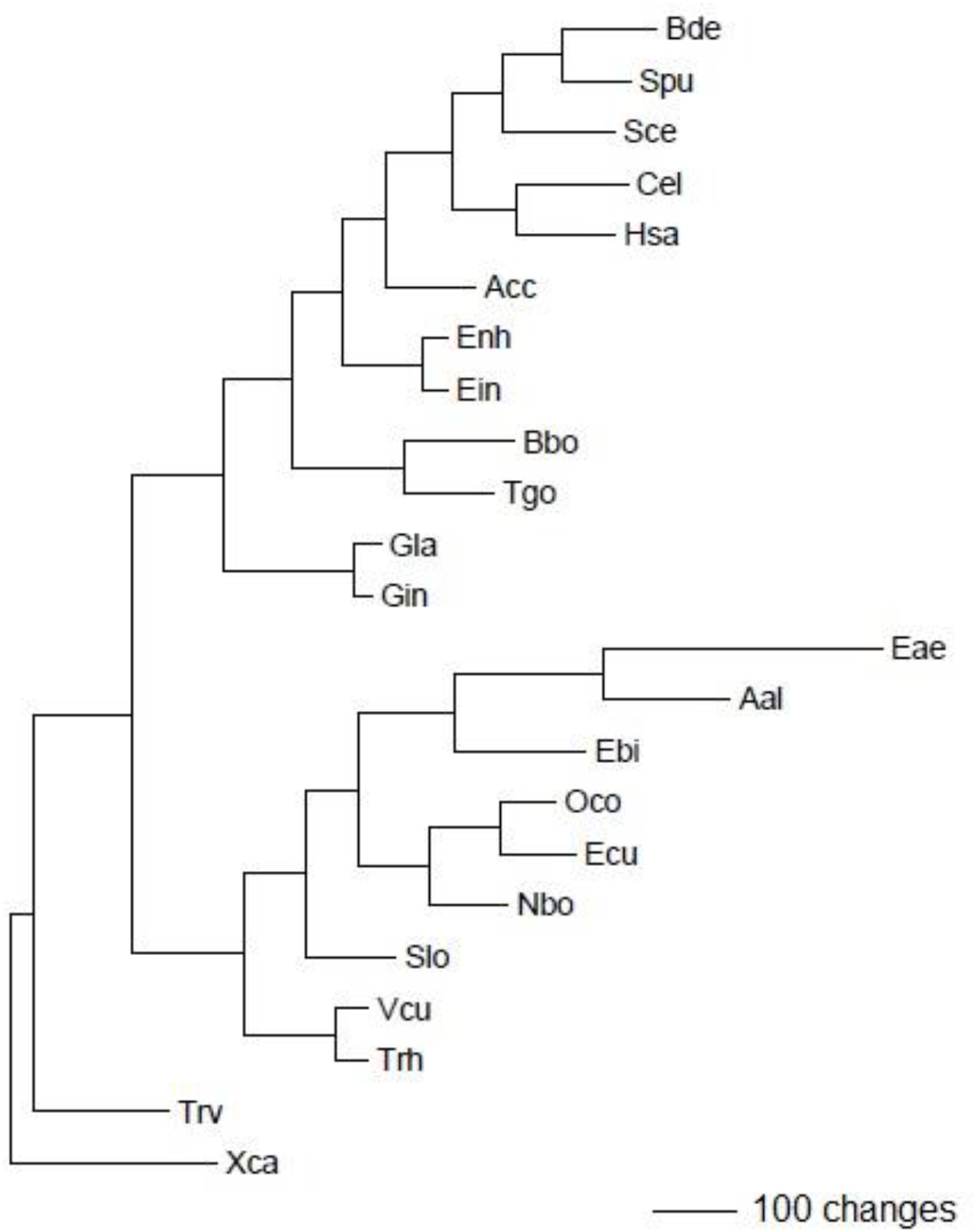
Phylogenetic tree of eukaryotic VARS sequences.

Accordingly, although the degeneration theory of AMI formation has been widely employed to explain the origin of AMIs, these explanations were saddled unavoidably with directional ambiguity. The acquisition of hydrogenase and PFO genes by the AMIs from the Tte-Hth-Ssp group through ectosymbiosis (Figure 4) was incompatible with the origin of AMIs from degenerating Fungi; and *Giardia* evolved prior to the MTEs including Fungi in the VARS phylogeny (Figure 6). Furthermore, the degeneration theory was invalidated by two different multi-protein fungal phylogenies, where Microsporidia represented either the earliest diverging clade of sequenced Fungi^49^, or the lowest branch on the fungal tree together with *Rozella*^50^, instead of the end products of other Fungi. It was necessary, however, to determine whether Microsporidia were older or younger than other AMIs such as *Giardia* and *Trichomonas*.

### Ectosymbiosis-based mechanism of eukaryogenesis

The heatmap in Figure 1 showed widespread influx of prokaryotic genes into the eukaryotes, which would include genes bequeathed to the eukaryotes by Abo as Archaeal Parent, and genes derived from archaea and bacteria through ecto- and endo-symbioses. Among the ectosymbiosis-derived genes, the Eco genes were outstanding for their continual adoption by both AMIs and MTEs, as indicated by the prominence of Eco genes in the DNA apparati of a variety of eukaryotes (Figure 2). On this basis, the ratio between the Abo-derived and Eco-derived genes in a eukaryote genome would provide a measure of how far the eukaryote had evolved from Abo. As shown in Figure 7, Microsporidia displayed the highest Abo/Eco ratios and therefore the oldest ages among the eukaryotes including Gla and Trv. The exceptionally low ratio displayed by Nbo among the Microsporidia corresponded to a strong Eco-to-Nbo gene transfer evident on the heatmap in Figure 1. In turn, the Gla ratio was greater than that of Trv, which in turn exceeded the MTE ratios. Thus the decreasing Abo/Eco ratio could be employed to track the order of emergence of different categories of eukaryotes, as outlined in the multistage eukaryogenesis mechanism in Figure 8.

**Figure 7.**
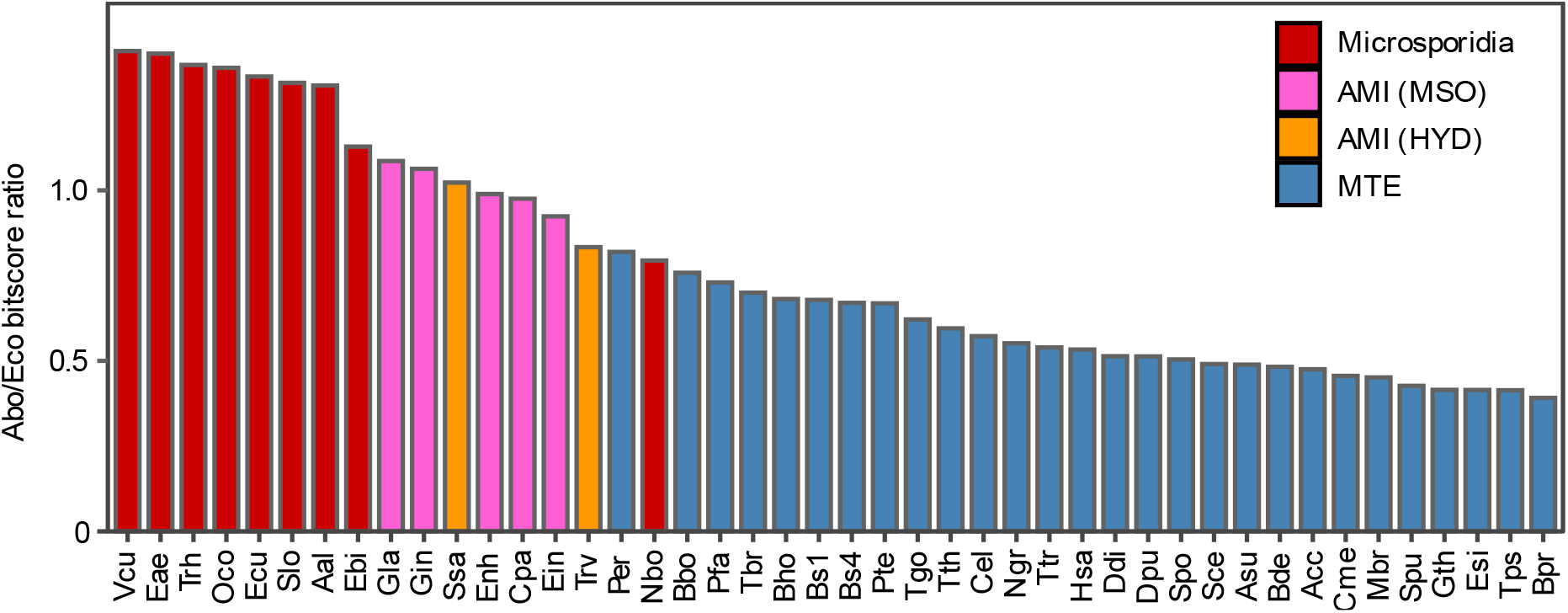
Ratios of eukaryotic similarity bitscores toward Abo relative to Eco. The eukaryotes (x-axis) are arranged in descending order of their Abo/Eco ratios. Columns of color-coded eukaryotes include Microsporidia; AMIs bearing mitosome (MSO); AMIs bearing hydrogenosome (HYD); and MTEs. See Supplementary Table 3 for the numerical ratios.

**Figure 8.**
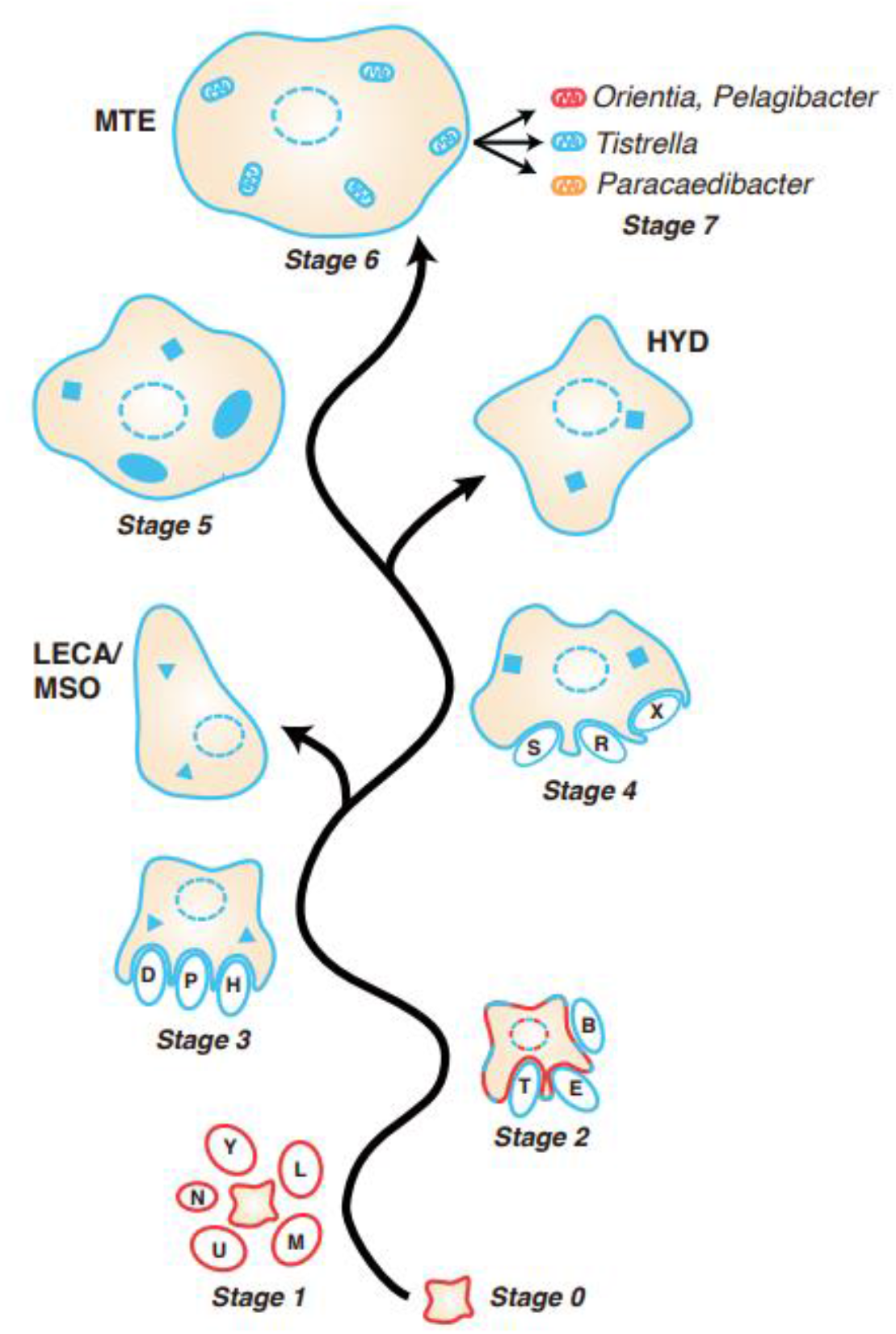
Ectosymbiotic mechanism of eukaryogenesis. Stages 0-7 are developmental stages of the Archaeal Parent lineage. Archaeal membranes are colored red, and bacterial membranes colored blue. Ellipses enclosed by dashed line represent cell nucleus; solid triangles mitosomes, solid squares hydrogenosomes, solid elipses endosymbionts, and elipses with cristae mitochondria. The letters Y, L, M, U, N, T, E, B, D, P, H, S, R and X stand for *Prometheoarchaeum, Lokiarchaeum, Methanosuratus, Pyrococcus*, DPANN, *Thermoanaerobacter, E. coli, B. subtilis, Paracoccus, Pelobacter, Hungateiclostridium, Tistrella, Ralstonia* and *Xanthomonas* respectively, representing just a small fraction of the archaea and bacteria taking part in eukaryogenesis as ectosymbiotic gene donors to the eukaryotes.

In this mechanism, Abo launched eukaryogenesis at Stage 0. At Stage 1, genes were recruited from archaeal and bacterial ectosymbionts, and the ether-lipid membranes of Abo were gradually replaced by ester lipids, and Abo underwent maturation as Archaeal Parent under the stimulus of mainly archaeal ectosymbionts. At Stage 2, the continual influx of exogenous genes brought about the formation of cell nucleus, marking the birth of LECA. Stages 3 and 4 witnessed the accelerated uptake and adoption of bacterial genes giving rise to the mitosome and hydrogenosome, and continual expansion of host cell volume as in the case of attainment of giant cell size by *Thaumarchaeota* through association with a gammaproteobacterial ectosymbiont^51^. The expanded cell volume enabled the capture and accomodation of proteobacterial ectosymbionts via phagocytosis^52^ or enwrapment with cytoplasmic membranes^53,54^ etc., turning them into endosymbionts and subsequently mitochondria at Stage 5, and finally mitochondria with different lineages of alphaproteobacteria mtDNA at Stages 6 and 7. Because the formation of cell nucleus preceded the capture of alphaproteobacterial endosymbionts, this mechanism represented an ectosymbiosis-based mechanism of eukaryogenesis^55^.

## DISCUSSION

Eukaryotes are known to contain a series of ESPs that are fundamental to all eukaryotes, and the Asgard archaeons were major sources of the ESPs in Gla, amounting to 39% of total ESPs with Tho, Odi, and Lok as the foremost contributors^8^, even though the Asgards might not have bridged the gap between prokaryotes and eukaryotes^56^. However, because there was no coalescence of the diverse Asgard ESP contributions through a single Asgard donor, Abo became the Archaeal Parent based on several criteria. First, the donations made by Abo toward the DNA apparati of AMIs, Fungi and *C. elegans* surpassed those made by any Asgard (Figure 2). Secondly, the total similarity hits exhibited by the Abo genes toward either Gla or Trv outnumbered those exhibited by other archaea tested^8^. Thirdly, Abo and Tvo were first and second among archaea in the strength of their AGA phenotype respectively, which would expedite the recruitment of exogenous genes by an Abo or Tvo Archaeal Parent and its offspring. Fourthly, Abo and Tvo were closely related^57^, but there were far fewer Tvo genes than Abo genes in the AMIs on account of the Archaeal Parenthood of Abo (Figure 1). In addition, the usefulness of the Abo/Eco ratio as a measure of the evolutionary distance traversed by any eukaryote from Abo was also indicative of the Archaeal Parent role of Abo.

The highest Abo/Eco ratios of Microsporidia among eukaryotes constituted key evidence for a Microsporidia-proximal LECA. In addition, the numerous top bitscores received by the microsporidian Eae, Vcu and Ecu COG markers in their DNA apparati from Abo attested to their ancient origins (Figure 2). As well, within the non-microsporidian division of the VARS tree in Figure 6, the *Giardia* species Gla and Gin, with Abo/Eco ratios smaller than those of Microsporidia, were lower branching than the MTEs including Fungi. The fungal phylogenies likewise indicate that Microsporidia were primitive Fungi rather than the derivatives of other Fungi49,50.

Between different Microsporidia, there were such divergence and high evolutionary rates that the utility of their sequence-based phylogenies has been questioned^58^. However, Microsporidia developed hexokinases bearing secretion signal sequences, and an elaborate extrusion device consisting of a coiled polar tube with an anchoring disc for their spores; and there are 150 genera and more than 1,200 species of Microsporidia capable of infecting virtually all animal phyla accompanied by rapid proliferation within the varied hosts^59^. They can even direct host biology to the formation of cyst-like xenoma that provided high concentrations of energy and nutrients to support massive growth of the microsporidian parasites^60^. Such vitality suggests that the divergence and rapid evolution of Microsporidia could be due to their adaptations to widely different hosts more than intrinsic genomic instability. This possibility was supported strongly by conserved aspects of microsporidian molecular biology, not only in the relative uniformity of their elevated Abo/Eco ratios but also in the conserved amino acid sequences of their VARS in the vicinity of the KMSK motif in line with other eukaryotes (Figure 5), and the clustering of all the Microsporidia within a separate division on the eukaryotic VARS tree (Figure 6).

In the ectosymbiosis-based mechanism (Figure 8), the proteome of Abo at Stage 0 contained 1,500 proteins, while a mitosome-containing eukaryote at Stage 3 exemplified by *Giardia* contained 5,000 proteins. This rate of proteome expansion would likely exceed the capability of HGT, but not ectosymbiosis especially when Abo the Archaeal Parent was equipped with top-ranked AGA activity. Evolutionary events mediated by ectosymbiosis in the present study would include the recruitment of a variety of archaeal and bacterial genes into the DNA apparati of eukaryotes (Figure 2); the appearance of multiple alphaproteobacterial gene segments within the same mtDNAs (Figure 3); the influx of genes for hydrogenase and PFO into the AMIs (Figure 4); the entry of a VARS with a 37-aa insert from Xca into the eukaryotes (Figure 5); and the ectosymbiont-assisted development of cell nucleus, mitosome and hydrogenosome (Figure 8). These diverse ectosymbioses were in accord with the widespread medium-intensity gene transfers indicated in the heatmap (Figure 1), as well as the historical utilization of different groups of bacterial genes by different higher taxa of archaea^61^.

## CONCLUSION

The three-domain structure of life proposed by Woese et al^1^ represents the core of biology. To understand more fully the functional significance of the separation of the three domains, elucidation of their origins becomes essential. In this regard, the identification by Xue et al^62^ of a LUCA proximal to *Methanopyrus kandleri* (Mka), a resident of deep-sea hydrothermal vents, has been confirmed by the top VARS-IARS bitscore of Mka among 5,000 species of organisms^8^; the invention of wobble rules by Mka in using uniformly the GNN and UNN anticodon duo to decode the four codons in all family tetracodon boxes, and employment of tRNAs(Ser) from adjacent sequence space to translate the non-contiguous UCN and AGY codons of Ser^63^; the hydrothermal vent-like habitat of LUCA^64^; and the oldest age of the *Methanopyrus* lineage among archaea dating back to 2.8 Gya^65^. These findings verified the prediction that such vents represented the birthplace of life on Earth^66,67^, which has turned the search for possible hydrothermal vents on Mars and other planets into a priority of exobiological exploration, and focused the search for an origin of eukaryotes within the Archaea.

Recently, evidence based on the use of dark fermentation by both *Thermococci* and primitive *Clostridia* for the production of hydrogen, and the sister-clade relationship between these two groups on the VARS phylogenetic tree has revealed a *Thermococci*-to-*Clostridia* evolutionary pathway for the emergence of Bacteria from Archaea^68^. As suggested by Nierhaus^69^, this emergence was driven by bacterial advances led by the acquisition of elongation factor 4 (EF4/LepA). EF4/LepA, highly conserved in bacteria but absent from archaea, catalyzed ribosomal back-translocation and remobilization of stalled ribosomes, thereby enhancing protein synthesis five-fold under conditions of high intracellular magnesium ion or low temperature^70^. Therefore it could increase the survival rates of bacterial derivatives of archaea compared to the archaea themselves in crossing the ice-cold seawater surrounding their original hydrothermal vent habitats to spread to the mesothermal zones.

While the origin of Archaea shaped an interface between the biological and inorganic worlds, and the origin of Bacteria from Archaea consisted of mainly vertical transmission of genes accompanied by extensive mutations and possibly some influx of exogenous genes, present evidence suggests that Eukarya depended on a completely different kind of origin that deployed a continual stream of ectosymbioses and a small number of endosymbioses. On account of the small size of Abo (1.4Mb genome), MTEs with upfront mitochondria generated by proteobacterial endosymbionts were extremely difficult to achieve owing to spatial constraint. As a result, the development of Abo as Archaeal Parent had to rely at first on prokaryotic genes recruited through ectosymbiosis. This enforced reliance presented a remarkable opportunity for the eukaryotes. Instead of extracting the genes from a handful of endosymbionts and becoming just highly capable archaeons, Abo and its offspring were free to pick and choose any number or variety of ectosymbiotically transmitted genes for adoption from a vast pool of prokaryotic genes equal to the accumulated innovations from eons of archaeal and bacterial evolution. Consequently, they could only enrich themselves with genes, mitosomes and hydrogenosomes until they were large enough in size to capture endosymbionts like alphaproteobacteria and cyanobacteria, and transform them into power-house organelles to drive their unending expansion. Without the immense advantages of ectosymbiosis, one can only ponder how many more billions of years it would take the Eukarya to attain such accomplishments as language and use of tools.

## Data availability

All data generated or analyzed during this study are included in this published article and its supplementary information files. The following are available online. Supplementary Table 1: Descriptions of species analyzed. Supplementary Table 2: Inter-proteome similarity bitscores between eukaryotes and prokaryotes. Supplementary Table 3: Abo/Eco ratios for eukaryotic proteomes toward Abo relative to Eco. Supplementary Figure 1: Organisms with or without a 37-aa insert in VARS downstream of the KMSK motif.

## Acknowledgments

This research was funded by grant number 5112-703-0110D-42002 from the University Grants Council of Hong Kong to Applied Genomics Center of Hong Kong University of Science and Technology.

## Author contributions

Conceptualization, J.T.W. and H.X.; data analysis, C.K.C. and X.L.; writing, J.T.W., C.K.C., X.L. and H.X.. All authors have read and agreed to the published version of the manuscript.

## References

1. Martin WF, Garg S, Zimorski V. Endosymbiotic theories for eukaryote origin. Phil Trans Roy Soc London B Biol Sci. 2015; 370: 20140330. doi: 10.1098/rstb.2014.0330.

2. Searcy DG, Stein DB, Searcy KB. A mycoplasma-like archaebacterium possibly related to the nucleus and cytoplasms of eukaryotic cells. Ann N Y Acad Sci. 1981;361:312–324. doi:10.1111/j.1749-6632.1981.tb46527.x.

3. Cox CJ, Foster PG, Hirt RP, Harris SR, Embley TM. The archaebacterial origin of eukaryotes. Proc Natl Acad Sci U S A. 2008;105(51):20356–20361. doi:10.1073/pnas.0810647105.

4. Woese CR, Kandler O, Wheelis ML. Towards a natural system of organisms: proposal for the domains Archaea, Bacteria, and Eucarya. Proc Natl Acad Sci U S A. 1990;87(12):4576–4579. doi:10.1073/pnas.87.12.4576.

5. Margulis L, Dolan MF, Guerrero R. The chimeric eukaryote: origin of the nucleus from the karyomastigont in amitochondriate protists. Proc Natl Acad Sci U S A. 2000;97(13):6954–6959. doi:10.1073/pnas.97.13.6954.

6. Cavalier-Smith T. The origin of eukaryotic and archaebacterial cells. Ann N Y Acad Sci. 1987;503:17–54. doi:10.1111/j.1749-6632.1987.tb40596.x.

7. Martin W, Müller M. The hydrogen hypothesis for the first eukaryote. Nature. 1998;392(6671):37–41. doi:10.1038/32096.

8. Long X, Xue H, Wong JTF. Descent of bacteria and eukarya from an archaeal root of life. Evol Bioinform Online. 2020;16:1176934320908267. Published 2020 Jun 23. doi:10.1177/1176934320908267.

9. Hartman H, Fedorov A. The origin of the eukaryotic cell: a genomic investigation. Proc Natl Acad Sci U S A. 2002;99(3):1420–1425. doi:10.1073/pnas.032658599.

10. Eme L, Spang A, Lombard J, Stairs CW, Ettema TJG. Archaea and the origin of eukaryotes. Nat Rev Microbiol. 2018;16(2):120. doi:10.1038/nrmicro.2017.154.

11. Raymann K, Brochier-Armanet C, Gribaldo S. The two-domain tree of life is linked to a new root for the Archaea. Proc Natl Acad Sci U S A. 2015;112(21):6670–6675. doi:10.1073/pnas.1420858112.

12. Cotton JA, McInerney JO. Eukaryotic genes of archaebacterial origin are more important than the more numerous eubacterial genes, irrespective of function. Proc Natl Acad Sci U S A. 2010;107(40):17252–17255. doi:10.1073/pnas.1000265107.

13. Vossbrinck CR, Maddox JV, Friedman S, Debrunner-Vossbrinck BA, Woese CR. Ribosomal RNA sequence suggests microsporidia are extremely ancient eukaryotes. Nature. 1987;326(6111):411–414. doi:10.1038/326411a0.

14. Sogin ML, Silberman JD. Evolution of the protists and protistan parasites from the perspective of molecular systematics. Int J Parasitol. 1998;28(1):11–20. doi:10.1016/s0020-7519(97)00181-1.

15. Kamaishi T, Hashimoto T, Nakamura Y, et al. Protein phylogeny of translation elongation factor EF-1 alpha suggests microsporidians are extremely ancient eukaryotes. J Mol Evol. 1996;42(2):257–263. doi:10.1007/BF02198852.

16. Pittis AA, Gabaldón T. Late acquisition of mitochondria by a host with chimaeric prokaryotic ancestry. Nature. 2016;531(7592):101–104. doi:10.1038/nature16941.

17. Horner DS, Hirt RP, Kilvington S, Lloyd D, Embley TM. Molecular data suggest an early acquisition of the mitochondrion endosymbiont. Proc Biol Sci. 1996;263(1373):1053–1059. doi:10.1098/rspb.1996.0155.

18. Roger AJ. Reconstructing early events in eukaryotic evolution. Am Nat. 1999;154(S4):S146–S163. doi:10.1086/303290.

19. Van de Peer Y, Ben Ali A, Meyer A. Microsporidia: accumulating molecular evidence that a group of amitochondriate and suspectedly primitive eukaryotes are just curious fungi. Gene. 2000;246(1-2):1–8. doi:10.1016/s0378-1119(00)00063-9.

20. Arisue N, Sánchez LB, Weiss LM, Müller M, Hashimoto T. Mitochondrial-type hsp70 genes of the amitochondriate protists, Giardia intestinalis, Entamoeba histolytica and two microsporidians. Parasitol Int. 2002;51(1):9–16. doi:10.1016/s1383-5769(01)00093-9.

21. Degli Esposti M. Late Mitochondrial acquisition, Really? Genome Biol Evol. 2016;8(6):2031–2035. doi:10.1093/gbe/evw130.

22. Martin WF, Roettger M, Ku C, Garg SG, Nelson-Sathi S, Landan G. Late mitochondrial origin is an artifact. Genome Biol Evol. 2017;9(2):373–379. doi:10.1093/gbe/evx027.

23. Gray MW. The pre-endosymbiont hypothesis: a new perspective on the origin and evolution of mitochondria. Cold Spring Harb Perspect Biol. 2014;6(3):a016097. Published 2014 Mar 1. doi:10.1101/cshperspect.a016097.

24. Clark K, Karsch-Mizrachi I, Lipman DJ, Ostell J, W. Sayers E, et al. GenBank. Nucleic Acids Res. 2016; 44 D67–72. doi: 10.1093/nar/gkv127.

25. O’Leary NA, Wright MW, Brister JR, Ciufo S, Haddad D, et al. Reference sequence (RefSeq) database at NCBI: current status, taxonomic expansion, and functional annotation. Nucl Acid Res. 2016; 44: D733–745. doi: 10.1093/nar/gkv1189.

26. Camacho C, Coulouris G, Avagyan V, Ma N, Papadopoulos J, Bealer K, Madden TL. BLAST+: architecture and applications. BMC Bioinform. 2009; 10: 421. doi: 10.1186/1471-2105-10-421.

27. Enderlin CS, Meeks JC. Pure culture and reconstitution of the Anthoceros-Nostoc symbiotic association. Planta. 1983; 158: 157–165. Doi: 10.1007/BF00397709.

28. Hétérier V, David B, De Ridder C, Rigaud T. Ectosymbiosis is a critical factor in the local benthic biodiversity of the Antarctic deep sea. Mar Ecol Prog Ser. Published online 2008. doi:10.3354/meps07487.

29. Noda S, Ohkuma M, Yamada A, Hongoh Y, Kudo T. Phylogenetic position and in situ identification of ectosymbiotic spirochetes on protists in the termite gut. Appl Environ Microbiol. 2003;69(1):625–633. doi:10.1128/AEM.69.1.625-633.2003.

30. Golyshina OV, Toshchakov SV, Makarova KS, et al. ‘ARMAN’ archaea depend on association with euryarchaeal host in culture and in situ. Nat Commun. 2017;8(1):60. Published 2017 Jul 5. doi:10.1038/s41467-017-00104-7.

31. Bauermeister J, Ramette A, Dattagupta S. Repeatedly evolved host-specific ectosymbioses between sulfur-oxidizing bacteria and amphipods living in a cave ecosystem. PLoS One. 2012;7(11):e50254. doi:10.1371/journal.pone.0050254.

32. Esser C, Ahmadinejad N, Wiegand C, Rotte C, Sebastiani F, Gelius-Dietrich G, Henze K, Kretschmann E, Richly E, Leister D, et al. A genome phylogeny for mitochondria among alpha-proteobacteria and a predominantly eubacterial ancestry of yeast nuclear genes. Mol Biol Evol. 2004; 21: 1643–1660. doi: 10.1093/molbev/msh160.

33. Rochette NC, Brochier-Armanet C, Gouy M. Phylogenomic test of the hypotheses for the evolutionary origin of eukaryotes. Mol Biol Evol. 2014; 31: 832–845. doi: 10.1093/molbev/mst27.

34. Tatusov RL, Galperin MY, Natale DA, Koonin EV. The COG database: a tool for genome-scale analysis of protein functions and evolution. Nucleic Acids Res. 2000;28(1):33–36. doi:10.1093/nar/28.1.33.

35. Andersson SG, Kurland CG. Origins of mitochondria and hydrogenosomes. Curr Opin Microbiol. 1999;2(5):535–541. doi:10.1016/s1369-5274(99)00013-2.

36. Abhishek A, Bavishi A, Bavishi A, Choudhary M. Bacterial genome chimaerism and the origin of mitochondria. Can J Microbiol. 2011;57(1):49–61. doi:10.1139/w10-099.

37. Wu B, Buljic A, Hao W. Extensive horizontal transfer and homologous recombination generate highly chimeric mitochondrial genomes in Yeast. Mol Biol Evol. 2015;32(10):2559–2570. doi:10.1093/molbev/msv127.

38. Ku C, Nelson-Sathi S, Roettger M, Garg S, Hazkani-Covo E, Martin WF. Endosymbiotic gene transfer from prokaryotic pangenomes: Inherited chimerism in eukaryotes. Proc Natl Acad Sci U S A. 2015;112(33):10139–10146. doi:10.1073/pnas.1421385112.

39. Martijn J, Vosseberg J, Guy L, Offre P, Ettema TJG. Deep mitochondrial origin outside the sampled alphaproteobacteria. Nature. 2018;557(7703):101–105. doi:10.1038/s41586-018-0059-5.

40. Whatley JM, John P, Whatley FR. From extracellular to intracellular: the establishment of mitochondria and chloroplasts. Proc R Soc Lond B Biol Sci. 1979;204(1155):165–187. doi:10.1098/rspb.1979.0020.

41. Horner DS, Hirt RP, Embley TM. A single eubacterial origin of eukaryotic pyruvate: ferredoxin oxidoreductase genes: implications for the evolution of anaerobic eukaryotes. Mol Biol Evol. 1999;16(9):1280–1291. doi:10.1093/oxfordjournals.molbev.a026218.

42. Tielens AG, Rotte C, van Hellemond JJ, Martin W. Mitochondria as we don’t know them. Trends Biochem Sci. 2002;27(11):564–572. doi:10.1016/s0968-0004(02)02193-x.

43. Hirt RP, Logsdon JM Jr, Healy B, Dorey MW, Doolittle WF, Embley TM. Microsporidia are related to Fungi: evidence from the largest subunit of RNA polymerase II and other proteins. Proc Natl Acad Sci U S A. 1999;96(2):580–585. doi:10.1073/pnas.96.2.580.

44. Keeling PJ. Congruent evidence from α-tubulin and β-tubulin gene phylogenies for a zygomycete origin of Microsporidia. Fungal Genet Biol. 2003: 298–309. doi:10.1016/S1087-1845(02)00537-6.

45. Roger AJ, Svärd SG, Tovar J, Clark CG, Smith MW, et al. A mitochondrial-like chaperonin 60 gene in Giardia lamblia: Evidence that diplomonads once harbored an endosymbiont related to the progenitor of mitochondria. Proc Nat Acad Sci USA 1998; 95: 229–234. doi:10.1073/pnas.95.1.229.

46. Blanchard JL, Lynch M. Organellar genes: why do they end up in the nucleus?. Trends Genet. 2000;16(7):315–320. doi:10.1016/s0168-9525(00)02053-9.

47. Marienfeld J, Unseld M, Brennicke A. The mitochondrial genome of Arabidopsis is composed of both native and immigrant information. Trends Plant Sci. 1999;4(12):495–502. doi:10.1016/s1360-1385(99)01502-2.

48. Hashimoto T, Sánchez LB, Shirakura T, Müller M, Hasegawa M. Secondary absence of mitochondria in Giardia lamblia and Trichomonas vaginalis revealed by valyl-tRNA synthetase phylogeny. Proc Natl Acad Sci U S A. 1998;95(12):6860–6865. doi:10.1073/pnas.95.12.6860.

49. Capella-Gutiérrez S, Marcet-Houben M, Gabaldón T. Phylogenomics supports microsporidia as the earliest diverging clade of sequenced fungi. BMC Biol. 2012;10:47. Published 2012 May 31. doi:10.1186/1741-7007-10-47.

50. James TY, Kauff F, Schoch CL, et al. Reconstructing the early evolution of Fungi using a six-gene phylogeny. Nature. 2006;443(7113):818–822. doi:10.1038/nature05110.

51. Muller F, Brissac T, Le Bris N, Felbeck H, Gros O. First description of giant Archaea (Thaumarchaeota) associated with putative bacterial ectosymbionts in a sulfidic marine habitat. Environ Microbiol. 2010;12(8):2371–2383. doi:10.1111/j.1462-2920.2010.02309.x.

52. Yutin N, Wolf MY, Wolf YI, Koonin EV. The origins of phagocytosis and eukaryogenesis. Biol Direct. 2009;4:9. Published 2009 Feb 26. doi:10.1186/1745-6150-4-9.

53. Baum DA, Baum B. An inside-out origin for the eukaryotic cell. BMC Biol. 2014;12:76. Published 2014 Oct 28. doi:10.1186/s12915-014-0076-2.

54. Imachi H, Nobu MK, Nakahara N, et al. Isolation of an archaeon at the prokaryote-eukaryote interface. Nature. 2020;577(7791):519–525. doi:10.1038/s41586-019-1916-6.

55. Wong JT-F, Chan C-K, Long X, Xue H. An ectosymbiosis-based mechanism of eukaryogenesis. bioRxiv. https://doi.org/10.1101/2022.10.20.512992

56. Da Cunha V, Gaia M, Gadelle D, Nasir A, Forterre P. Lokiarchaea are close relatives of Euryarchaeota, not bridging the gap between prokaryotes and eukaryotes. PLoS Genet. 2017;13(6):e1006810. Published 2017 Jun 12. doi:10.1371/journal.pgen.1006810.

57. Reysenbach AL, Liu Y, Banta AB, et al. A ubiquitous thermoacidophilic archaeon from deep-sea hydrothermal vents. Nature. 2006;442(7101):444–447. doi:10.1038/nature04921.

58. Corradi N, Keeling PJ. Microsporidia: a journey through radical taxonomical revisions. Fungal Biol Rev. Published online 2009. doi:10.1016/j.fbr.2009.05.001.

59. Cuomo CA, Desjardins CA, Bakowski MA, et al. Microsporidian genome analysis reveals evolutionary strategies for obligate intracellular growth. Genome Res. 2012;22(12):2478–2488. doi:10.1101/gr.142802.112.

60. Li T, Fang Z, He Q, Yu B, Zhou Z. Characterizing the xenoma of Varimorpha necatrix provides insights into the most efficient mode of Microsporidian proliferation. Front Cell Infect Microbiology 2021. Doi: 10.3389/fcimb.2021.699239.

61. Nelson-Sathi S, Sousa FL, Roettger M, et al. Origins of major archaeal clades correspond to gene acquisitions from bacteria. Nature. 2015;517(7532):77–80. doi:10.1038/nature13805

62. Xue H, Tong KL, Marck C, Grosjean H, Wong JTF. Transfer RNA paralogs: evidence for genetic code-amino acid biosynthesis coevolution and an archaeal root of life. Gene. 2003;310:59–66. doi:10.1016/s0378-1119(03)00552-3.

63. Wong JTF, Ng SK, Mat WK, Hu T, Xue H. Coevolution theory of the genetic code at age forty: pathway to translation and synthetic life. Life. 2016;6(1):12. Published 2016 Mar 16. doi:10.3390/life6010012

64. Weiss MC, Preiner M, Xavier JC, Zimorski V, Martin WF. The last universal common ancestor between ancient Earth chemistry and the onset of genetics. PLoS Genet. Published online 2018. doi:10.1371/journal.pgen.1007518

65. Blank CE. Low rates of lateral gene transfer among metabolic genes define the evolving biogeochemical niches of archaea through deep time. Archaea 2012; 2012: 23. Doi: 10.1155/2012/843539.

66. Corliss JB, Dymond J, Gordon LI, et al. Submarine thermal springs on the Galápagos Rift. Science. Published online 1979. doi:10.1126/science.203.4385.1073

67. Baross JA, Hoffman SE. Submarine hydrothermal vents and associated gradient environments as sites for the origin and evolution of life. Orig Life Evol Biosph. Published online 1985. doi:10.1007/BF01808177

68. Wong TF, Chan CK, Xue H. Thermococcus-to-Clostridia pathway for the evolution of the Bacteria domain. bioRxiv. Preprint posted online October 21, 2022. https://doi.org/10.1101/2022.10.20.512992

69. Nierhaus KH. Cited in Wong JTF, Root of Life. In Prebiotic Evolution and Astrobiology; Wong, JTF., Lazcano, A., eds.; Landes Bioscience: Austin, TX, USA, 2009; pp. 120–144. doi:10.1201/9781498713986.

70. Pech M, Karim Z, Yamamoto H, Kitakawa M, Qin Y, Nierhaus KH. Elongation factor 4 (EF4/LepA) accelerates protein synthesis at increased Mg2+ concentrations. Proc Natl Acad Sci U S A. 2011;108(8):3199–3203. doi:10.1073/pnas.1012994108

