## Supplementary- Figure 1 for "An ectosymbiosis-based mechanism of eukaryogenesis"

**Supplementary Figure 1. Species or organisms with or without a 37-aa insert in VARS downstream of the KMSK motif. All sequences are numbered according to the Xca sequence which begins with residue 741 and ends with residue 816 in the alignment below. Notably, the algae Cme and Bpr among the eukaryote were devoid of the insert.**

ARCHAEA

Abo ILAPDGRPMHTSWGNV--VDPLEIID-------------------------------------EYGADALRFFAA

Afu VFGEDGRKMSKSLGNV--IVPEEVVE-------------------------------------KYGVDALRQWAA

Hei VVDSKGEKLSKSKGTD--VQPEKMIE-------------------------------------KYGGDAVRFYGA

Mac VLGPDGHKMSKSLGNV--ISPEEVTT-------------------------------------QYSADAFRQWGA

Mbo VLGEDGFKMSKSRGNV--IVPEDLVG-------------------------------------RYGADALRQWAA

Mco GLDPHGKAMHKSKGNI--VEPLPIVD-------------------------------------KYSADALRWWAA

Mja VFGEDGHKMSKSRGNV--VEPDEIIA-------------------------------------KYGADALRLWAS

Odi VLDEHGRAMHKSLGNI--VWVEPLLK-------------------------------------KYGADALRLFGC

Pfu VAGPDGRKMSKSYGNV--VSPEEVIP-------------------------------------KYGADALRLWTA

Psy IRDAKGQKISKSMENIEDYDPLKIIE-------------------------------------NVGADSLRYALI

Sso VLGPDGTRMSKSKGNV--VSPLDRVN-------------------------------------DFGADAIRMALL

Tho VVDENGETMSKSKGNS--PPPMPFVE-------------------------------------KYGADAMRMFGI

Tvo VFDMYGEKMSKSKGNI--VDIYAITD-------------------------------------KYGADALRFWAS

BACTERIA

Aba VRDGEGQKMSKSKGNV--LDPLDLIDGIDLESLVAKRTTGLMNPKDAAKIEKSTRKEFPEGINAYGTDAVRFTFC

Atu VRDKNGQKMSKSKGNV--IDPLELID-------------------------------------EYGADALRFTLA

Bja VRDEKGAKMSKSKGNV--IDPLNLID-------------------------------------EYGADALRFTLA

Cje VKDEQGRKMSKSLGNV--IDPNESIK-------------------------------------EYSADILRFTLA

Mtu IRDESGRKMSKSKGNV--IDPLDWVE-------------------------------------MFGADALRFTLA

Pde VRDEKGAKMSKSKGNV--IDPLTLID-------------------------------------EYGADALRFTLT

Pel VRDASGQKMSKSKGNV--IDPLTVID-------------------------------------EYGTDAFRFTLA

Rru VRDEKGQKMSKSKGNV--IDPLDMTD-------------------------------------QYGTDALRFTLI

Rso VRDSEGKKMSKSEGNT--LDPVDLIDGIALEPLLVKRTTGLRRPKDAPNVEKRTRKEFPDGIPAFGADALRFTFA

Ssp VRDSQGRKMSKSLGNG--IDPLDVID-------------------------------------KYGADALRFTLV

Syn VRDENGKKMSKSANNG--IDPLLLIN-------------------------------------KYGTDALRYTLI

Tis VRDEKGQKMSKSKGNV--IDPIDLID-------------------------------------KYGADAVRFTLL

Tte VRDALGRKMSKSLGNG--IDPLEVIE-------------------------------------KYGADTLRFTLV

Eco IRDDEGQKMSKSKGNV--IDPLDMVDGISLPELLEKRTGNMMQPQLADKIRKRTEKQFPNGIEPHGTDALRFTLA

Xca IRDAQGQKMSKSKGNV—LDPLDIIDGISIEDLVAKRTSGLMQPRMAEKIEKATRKEFPDGIIAHGADALRFTIA

MICROSPORIDIAN

Vcu VRDANGRKMSKSLGNVIDPLYVIEGIELDELAKSVT----STNLEPREVKTALEGQKKDFPMGIPRCGSDALRFT

Eae IRDAHGKKMSKSLGNVIDPLFIINGIKLSEMNNILKESHNNGYISNQELLRALDSQKKDFPRGVANCGADALRFA

Trh VRDANGRKMSKSLGNVIDPLYVIEGVQLDELAKSIT----ATNLDPKEIKAALEGQRKDFPMGIPRCGSDALRFT

Oco VRDAHGRKMSKSLGNVIDPIFVIDGCSLNELIATMK----SGNLDEKEVKVAEAALRKDFPNGIPRCGADALRFT

Ecu VRDAHGRKMSKSLGNVIDPIFVIDGCSLEKLISTMR----SGNLDEREVKRAEAVLRQDFPNGISRCGADALRFA

Slo VRDTHGRKMSKSLGNVIDPIFVIEGISLKGLNESI-----MTNLDKDEIKKAIEGQKKEYPNGIPQCGADALRFA

Aal IRDSMGRKMSKSLGNVIDPLFIINGCELKELNDSI-----SSTLSKKERDISLTYQKKTFPNGIKKCGADALRFC

Ebi VRDAHGKKMSKSLGNVIDPIFVIDGASQEELISKI-----SINVSNEEKKRAIASIKLDYPNGIPKCGADALRFA

Nbo VRDAHGRKMSKSLGNVIDPLFVIDGSSLENLIEVMK----SGNLALSEIKLAEKNLRKDFASGIAKCGADALRFT

OTHER EUKARYOTES

Gla VRDAHGAKMSKSKGNVVDPIDVIKGITLQEMGDKVR----ATNLPPKEIERALELQSKDFPIGIPECGTDALRFA

Gin VRDAHGAKMSKSKGNVVDPIDVIKGITLQEMGDKVR----ATNLPPKEIERALELQSKDFPIGIPECGTDALRFA

Cpa VRDSQGRKMSKSLGNVIDPIEIIEGISFDDLNKKLD----QGNLPLQEIKKSKENNLKDFPDGIPECGADALRIG

Enh VRDAQGRKMSKSLGNIIDPIDVIEGISLKGLNDKLY----TYNLPEKECVIAAEGQKKDFPNGIEECGTDAMRFA

Ein IRDAQGRKMSKSLGNVIDPIDVIEGISLEGLNEKLK----IYNLSEKEIAIATKGQQMNFPHGIEECGTDAMRFA

Ssa VRDSKGEKMSKSKGNVIDPLDCIFGISLKDLHARLR----EGNLSENEIKLAEKLQKAEFPAGIAQCGTDALRMA

Trv VRDAQGRKMSKSLGNVIDPRHVINGIELEDLVAEIE----NSTFDDKEKKIAIDGRKADFPNGIPQCGTDAMRLA

Tbr VRDKNGEKMSKSKGNVIDPLFIISGVSLEALHDTVR----SGNLDEKEVSRALKLQRETFPNGIPECGSDALRFG

Bbo VRDARGEKMSKSKGNVLDPLEVIEGATLDSLIDKIN----NSSLPQGEIKKAIVLQKQQFPQGIPACGTDALRLG

Pfa IRDSRGEKMSKSKGNVVDPLDIIDGISLNKLHEKLY----EGNLPEKEIKRAIELQKKEFPKGIPECGTDALRFG

Bho IRDKYGRKMSKSLGNVIDPLEIINGCDLESMLEKIR----HGNLDPAEVERASQGKRQDFPEGIPMCGTDALRFG

Pte IRDKDGKKMSKSLGNVIDPLEIIDGTSLENLKSKIY----EGNLSKDEVERAIKQKEEEFPNGIPECGGDALRFG

Tgo VRDAHGQKMSKSKGNVIDPLEVISGISLQDLQAKLH----KGNLPEKEIKRAEEVLKKEFPKGIEACGCDALRLG

Tth IRDSQGEKMSKSKGNVIDPLEIIDGCNLQTLIQKIQ----EGNLDKKEMNRAVQLKSKEYPEGFPECGSDALRYG

Ngr VRDKQGRKMSKSLGNVIDPIDMIKGTTFEDLKRGIE---KNTNITKQEMKKALQGVQQEFPNGIPQCGTDALRFT

Ddi VRDSHGRKMSKSLGNVIDPNDVIKGISLDELIAKLY----EGNLDSKEIEKATSGVKSDFPTGIAECGTDAMRFA

Dpu IRDSQGRKMSKSLGNVIDPLNVINGITLKELKDNVL----SSNLTDKEKSIATKGLDKEFPQGIPQCGTDSLRLS

Asu VRDSHGRKMSKSLGNVIDPLEVISGITLDQLVENLK----KGNLDPVELDRATLGLKQDYPEGITECGTDAMRFA

Acc VRDAHGRKMSKSLGNVVDPIDVTEGIRLTDMHQKLR----EGNLEASEVEKAIKGQQKDFPNGISECGTDAMRFA

Cme VRDANGRKMSKTLGNVTDPLEVI-----------------------------------------SKYGTDALRFT

Esi VRDKFGRKMSKTLGNVIDPLEVIYGCDLDTLHKKLE----VGNLPAKEMQKAKEGQKMDFPKGIPECGADALRFG

Tps VRDKDGRKMSKSLGNVIDPLEVINGCTLETLLEKLE----GGNLPPKEVARAKKDQEADFPEGIPECGSDALRFG

Bpr VRDEQGRKMSKSLGNVVDPLGVI-----------------------------------------GDVGCDALRFT

Ttr VRDKSGRKMSKSLGNVLDPADLIQGASLDKLLAALE----GGNLPAGERARASSDLKAAFPDGFPAFGTDALRYA

Spo VRDSEGRKMSKSLGNVIDPMDIINGVTLENMKKALL----EGNLPISEVHKSSKQMEKAFPNGIPAQGIDIFRYG

Sce VRDAQGRKMSKSLGNVIDPLDVITGIKLDDLHAKLL----QGNLDPREVEKAKIGQKESYPNGIPQCGTDAMRFA

Bde IRDAHGRKMSKSLGNVIDPIDVIEGVTLQLLQERLE----KGNLDPRELVRARDGQKKDFPNGIPECGTDALRFG

Spu VRDAHGRKMSKSLGNVIDPMDVINGISLPLLHKRLE----EGNLDAREIKKAQEGQKRDFPNGIPQCGTDAMRFA

Mbr IRDKEGRKMSKSLGNVVDPMDVRNGITLEDLHERLL----EGNLDPAELERAKEGQKRQFPDGIKECGVDALRFA

Cel IRDAHGRKMSKSLGNVIDPLDVIRGISLNDLQAQLL----GGNLDEKEIAVAKEGQARDYPDGIPECGVDALRFA

Has VRDAHGRKMSKSLGNVIDPLDVIYGISLQGLHNQLL----NSNLDPSEVEKAKEGQKADFPAGIPECGTDALRFG
